# Membrane progesterone receptors PAQR6 and PAQR8 in the ruminant uterus and ovary

**DOI:** 10.64898/2026.09.25.754262

**Authors:** Haoyu Yang, Yuki Yamamoto, Mayu Yamano, Yuta Matsuno, Kazuhiko Imakawa, Fumitaka Osakada, Hiroaki Ohno, Eiji Kondoh, Ikuo Kimura, Kentaro Nagaoka

## Abstract

Progesterone governs the establishment and maintenance of pregnancy in ruminants, acting through the nuclear progesterone receptor (PGR) and, potentially, through membrane progesterone receptors (mPRs) of the PAQR family. The cellular distribution of mPRs in ruminant reproductive tissues, however, remains undefined. We generated and validated rabbit polyclonal antibodies against the intracellular C-terminal peptides of bovine PAQR6 (mPRδ) and PAQR8 (mPRβ), and used them, with quantitative PCR of the uterus and conceptus, to map these receptors in the ovine uterus and conceptus and in the bovine ovary. Both antisera showed high ELISA titer against their immunizing peptides, negligible preimmune reactivity, and complete loss of tissue signal after peptide pre-absorption. By qPCR, *PAQR6*, *PAQR8* and *PGR* transcript levels did not change significantly across the peri-implantation period in either the conceptus or the endometrium; in contrast, the progesterone-responsive gene ZBTB16 increased markedly in the conceptus (up to ∼45-fold by day 21, p < 0.05 at days 17, 19 and 21), whereas FKBP5 showed a numerical increase without reaching statistical significance. Both genes remained relatively stable in the endometrium, indicating conceptus-specific changes in progesterone-responsive gene expression. At the protein level, PAQR6 and PAQR8 localized to the luminal and glandular epithelium of the endometrium across the cycle and early pregnancy, and to follicular (granulosa) and luteal cells of the ovary. Dual immunofluorescence in the ovine endometrium showed that both PAQR6 and PAQR8 are enriched at the apical surface of the luminal epithelium and are spatially distinct from the basolateral adherens-junction marker E-cadherin. These results provide a protein-level map of PAQR6 and PAQR8 in the ruminant reproductive tract, identify the apical epithelial surface as a site of receptor enrichment, and show that the endometrial epithelium retains membrane progesterone receptors while progesterone-responsive gene expression changes dynamically in the conceptus during the peri-implantation period. The validated ruminant-reactive antibodies provide a resource for future functional studies.

## INTRODUCTION

Progesterone is the central hormone of pregnancy in ruminants. Secreted by the corpus luteum, it acts on the endometrium to support conceptus elongation, regulate histotroph secretion from the uterine glands, and suppress the luteolytic pulses of prostaglandin F2α that would otherwise end the cycle (McCracken et al., 1999; Spencer and Bazer, 2002). In cyclic and pregnant sheep, progesterone progressively down-regulates the nuclear progesterone receptor (PGR) in the luminal and superficial glandular epithelium through continuous exposure, and this loss of epithelial PGR is a prerequisite for the onset of endometrial receptivity (Spencer and Bazer, 1995; Spencer et al., 2004). Once the epithelium has lost PGR, these genomic actions of progesterone are sustained mainly by the PGR-retaining stroma and deep glandular epithelium, which drive progesterone-responsive genes (e.g. *FKBP5*, Hubler and Scammell, 2004; *ZBTB16*/PLZF, Kommagani et al., 2016) and paracrine mediators central to peri-implantation biology.

Progesterone also elicits rapid, membrane-initiated responses that cannot be explained by the direct transcriptional action of nuclear PGR alone. These are attributed in part to membrane progesterone receptors (mPRs), a group of five seven-transmembrane proteins of the progestin and adipoQ receptor (PAQR) family: PAQR7 (mPRα), PAQR8 (mPRβ), PAQR5 (mPRγ), PAQR6 (mPRδ) and PAQR9 (mPRε) (Zhu et al., 2003b; Tang et al., 2005; Thomas, 2008). As members of the PAQR family they couple to G proteins and modulate second messengers within seconds to minutes, independently of the receptor acting as a nuclear transcription factor (Zhu et al., 2003a; Thomas, 2008). mPRs were first defined in fish oocytes, where they mediate progestin-induced meiotic maturation, and have since been implicated in the myometrium, oviduct and ovary of mammals (Zhu et al., 2003a; Dressing et al., 2011); in cattle, mPR transcripts have been reported in the corpus luteum and uterus (Kowalik et al., 2018; Kowalik et al., 2019). Their protein distribution in the ruminant uterus, conceptus and ovary, however, remains largely unknown.

Progress has been limited by the scarcity of validated antibodies that recognize ruminant PAQR proteins. Most commercial reagents are raised against human or rodent sequences and are not verified in cattle or sheep. To address this, we raised rabbit polyclonal antibodies against C-terminal intracellular peptides of bovine PAQR6 and PAQR8, the two subtypes we focus on here, and validated them by ELISA, preimmune-serum comparison and peptide pre-absorption. Using these antibodies together with quantitative PCR, we compared the expression of PAQR6, PAQR8 and the PGR genomic axis (PGR, ZBTB16, FKBP5) between the conceptus and the endometrium across the peri-implantation period, mapped PAQR6 and PAQR8 protein in the ovine endometrium and the bovine ovary, and located PAQR6 and PAQR8 relative to the epithelial cell membrane by co-staining with the basolateral adherens-junction marker E-cadherin.

## MATERIALS AND METHODS

### Animals and tissues

Tissues were collected from whiteface crossbred ewes maintained at the University of Tennessee (Knoxville, TN, USA), as part of collaborative work with Dr James D. Godkin. The protocol was reviewed and approved by the Animal Care Committees of the University of Tennessee, the University of Tokyo and Tokai University (approval number 20-017-23). Animal husbandry, estrous synchronization and tissue collection were performed as described previously (Imakawa et al., 2002). Conceptuses and endometria were collected across the estrous cycle and early pregnancy and frozen at −80 °C until use (Imakawa et al., 2006; Sakurai et al., 2012). Fixed, paraffin-embedded uteri and frozen conceptuses and endometria were transferred to the Laboratory of Animal Breeding, the University of Tokyo, and then to Tokai University (Kumamoto, Japan). Bovine ovaries were obtained from three animals at a local abattoir.

### Antibody generation

Rabbit polyclonal antibodies were raised against synthetic C-terminal intracellular peptides of bovine PAQR6 (residues 331–344, NH2-C+QGGSLERGTKAKLQ-COOH) and bovine PAQR8 (residues 341–354, NH2-C+LRQKIKARLTKKDS-COOH); the N-terminal cysteine was added for carrier conjugation. The corresponding ovine sequences differ by one to two residues (Fig. 1A), and the antibodies detected native protein in ovine tissue despite these mismatches. Peptide synthesis, KLH conjugation, rabbit immunization and antiserum collection were carried out as a custom antibody service by Eurofins Genomics (Tokyo, Japan). Antiserum titer against the immunizing peptide was determined by ELISA against a dilution series, with preimmune serum from the same animal as a control.

**Fig. 1.**
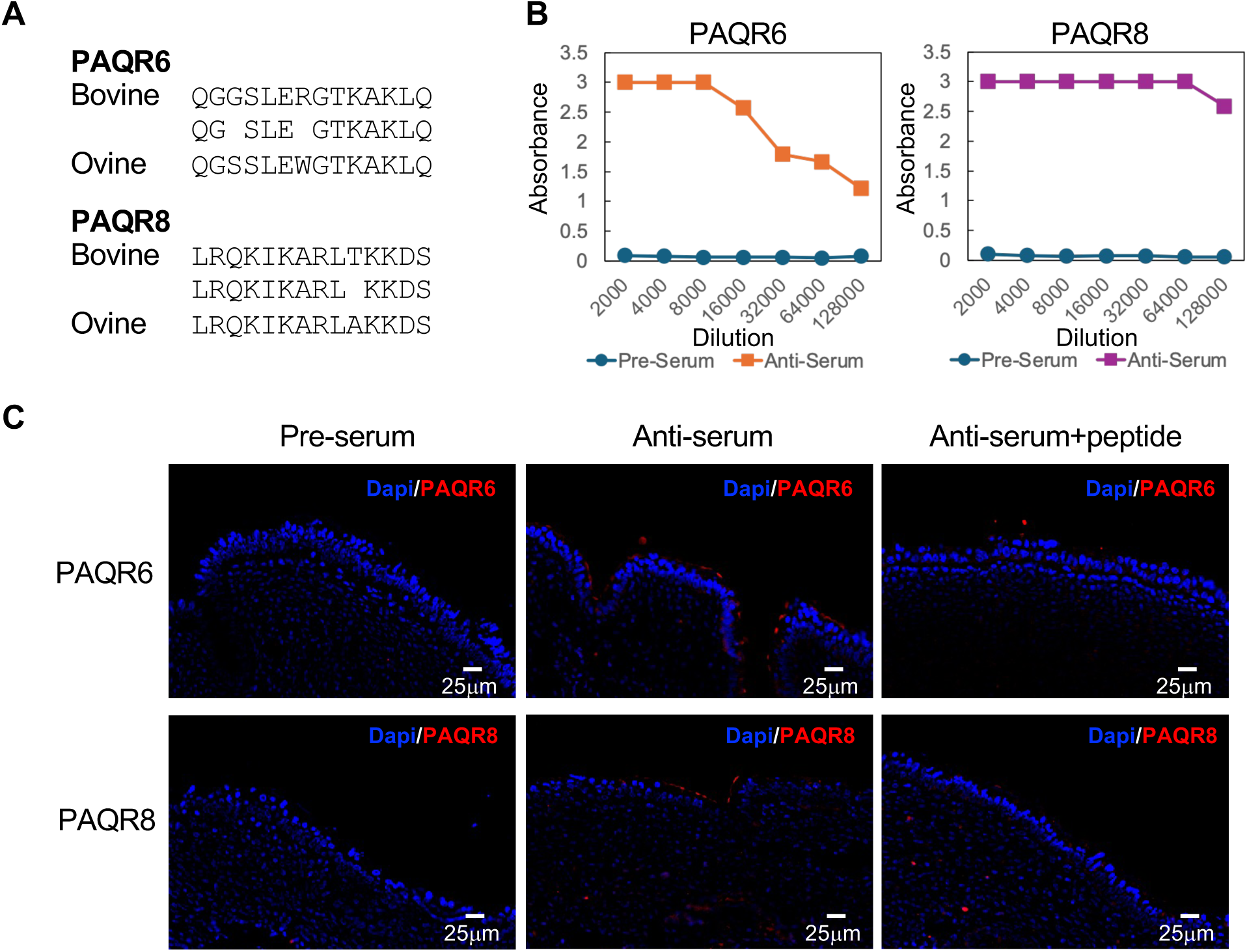
Generation and validation of antibodies against bovine PAQR6 and PAQR8. (A) Alignment of the immunizing bovine C-terminal peptides with the corresponding ovine sequences for PAQR6 (residues 331–344) and PAQR8 (residues 341–354); conserved residues are indicated. (B) ELISA titration of anti-PAQR6 and anti-PAQR8 antisera against their immunizing peptides across a dilution series, with matched preimmune (pre-) serum. (C) Immunofluorescence on ovine endometrium for PAQR6 (top) and PAQR8 (bottom) with preimmune serum, anti-serum, and anti-serum after pre-absorption with the corresponding peptide; nuclei counterstained with DAPI, all images at identical exposure. Peptide pre-absorption abolished specific signal. Scale bars, 25 µm.

### Quantitative PCR

Total RNA was extracted with ISOGEN II (Nippon Gene) and reverse-transcribed with the PrimeScript RT reagent Kit (Perfect Real Time; Takara Bio). Quantitative real-time PCR was performed with the KAPA SYBR FAST qPCR Kit (Roche, Cat. No. KK4602) on a 7500 Fast Real-Time PCR System (Applied Biosystems). Each 10 µL reaction contained 5 µL of 2× KAPA SYBR FAST qPCR Master Mix, 0.2 µL of 50× ROX Low reference dye, 2 µL of cDNA template, 0.2 µL each of forward and reverse primer, and 2.4 µL of nuclease-free water. Cycling consisted of initial denaturation at 95 °C for 2 min, followed by 40 cycles of 95 °C for 5 s and 60 °C for 30 s (annealing/extension). Transcripts of *PAQR6*, *PAQR8*, *PGR*, *ZBTB16* and *FKBP5* were quantified in the conceptus and the endometrium, with each sample run in technical duplicate (n = 3 biological replicates per group). Relative expression was calculated by the ΔΔCt method with *UXT* as the reference gene, normalized to day 15 (conceptus) or to cyclic day 15 (C15, endometrium). Primer sequences are listed in Supplementary Table S1.

### Immunofluorescence and co-localization

Paraffin sections underwent antigen retrieval in citrate buffer (pH 6.0) by microwaving (500 W, 15 min) and were blocked with 5% BSA in PBS. Sections were incubated overnight at 37 °C with anti-PAQR6 (1:200) or anti-PAQR8 (1:200), alone or together with mouse monoclonal anti-E-cadherin (BD Biosciences #610181; 1:100 in PBS containing 0.1% BSA). Bound antibodies were detected with Alexa Fluor 647-conjugated goat anti-rabbit IgG (Thermo Fisher Scientific #A-21244) and Alexa Fluor 488-conjugated goat anti-mouse IgG (Thermo Fisher Scientific #A-11001), both at 1:500, for 1 h at room temperature. Sections were mounted in DAPI-containing antifade medium (Fluoro-KEEPER Antifade Reagent, Non-Hardening Type with DAPI; Nacalai Tesque #12745-74). Specificity controls, run in parallel on adjacent sections at identical exposure, comprised preimmune serum, primary-antibody omission, and pre-absorption of each anti-PAQR antibody with its immunizing peptide (50 µg/ml for anti-PAQR6 and 100 µg/ml for anti-PAQR8; Fig. 1C); a mouse IgG2a isotype antibody (clone E5Y6Q; Cell Signaling Technology #61656) served as the negative control for anti-E-cadherin.

### Statistics

Group differences were tested by one-way ANOVA on ΔCt values followed by Dunnett’s test against the day-15 reference group; p < 0.05 was considered significant. Data are presented as mean ± SEM.

## RESULTS

### Generation and validation of anti-PAQR6 and anti-PAQR8 antibodies

The immunizing bovine peptides differ from the corresponding ovine sequences by one residue (PAQR8) or two residues (PAQR6) (Fig. 1A). Both antisera recognized their immunizing peptides at high titer by ELISA, the anti-PAQR8 antiserum remaining near saturation to beyond 1:64,000 and the anti-PAQR6 antiserum titrating over the same range, whereas the matched preimmune sera were unreactive throughout (Fig. 1B). In ovine endometrium, each antibody produced a defined signal that was abolished by pre-absorption with the corresponding peptide, and no signal was seen with preimmune serum (Fig. 1C). The two immunizing peptides share few residues, making cross-recognition between PAQR6 and PAQR8 unlikely. The anti-PAQR8 antiserum gave more background than anti-PAQR6 and required a higher peptide concentration for complete pre-absorption; with reference to the unstained-channel control, the specific signal was identified as the peptide-blockable component.

### PAQR6 and PAQR8 are expressed in the ovine conceptus and endometrium

We compared transcript levels of *PAQR6*, *PAQR8* and the *PGR* genomic axis between the conceptus (days 15-21 of pregnancy) and the endometrium (cyclic day 15 and days 15-21 of pregnancy) (Fig. 2). *PAQR6*, *PAQR8* and *PGR* did not change significantly across the peri-implantation period in either tissue (Fig. 2A, B). In contrast, the two progesterone-responsive genes showed different expression patterns between the conceptus and endometrium. In the conceptus, *ZBTB16* increased progressively to ∼45-fold by day 21 (p < 0.05 at days 17, 19 and 21), whereas *FKBP5* showed a numerical increase to ∼8-fold by day 21 that did not reach statistical significance. In the endometrium, neither gene showed a significant change across the stages examined (Fig. 2A, B). Thus, progesterone-responsive gene expression shows more pronounced day-dependent changes in the conceptus than in the maternal endometrium, even though membrane receptor transcripts are present in both.

**Fig. 2.**
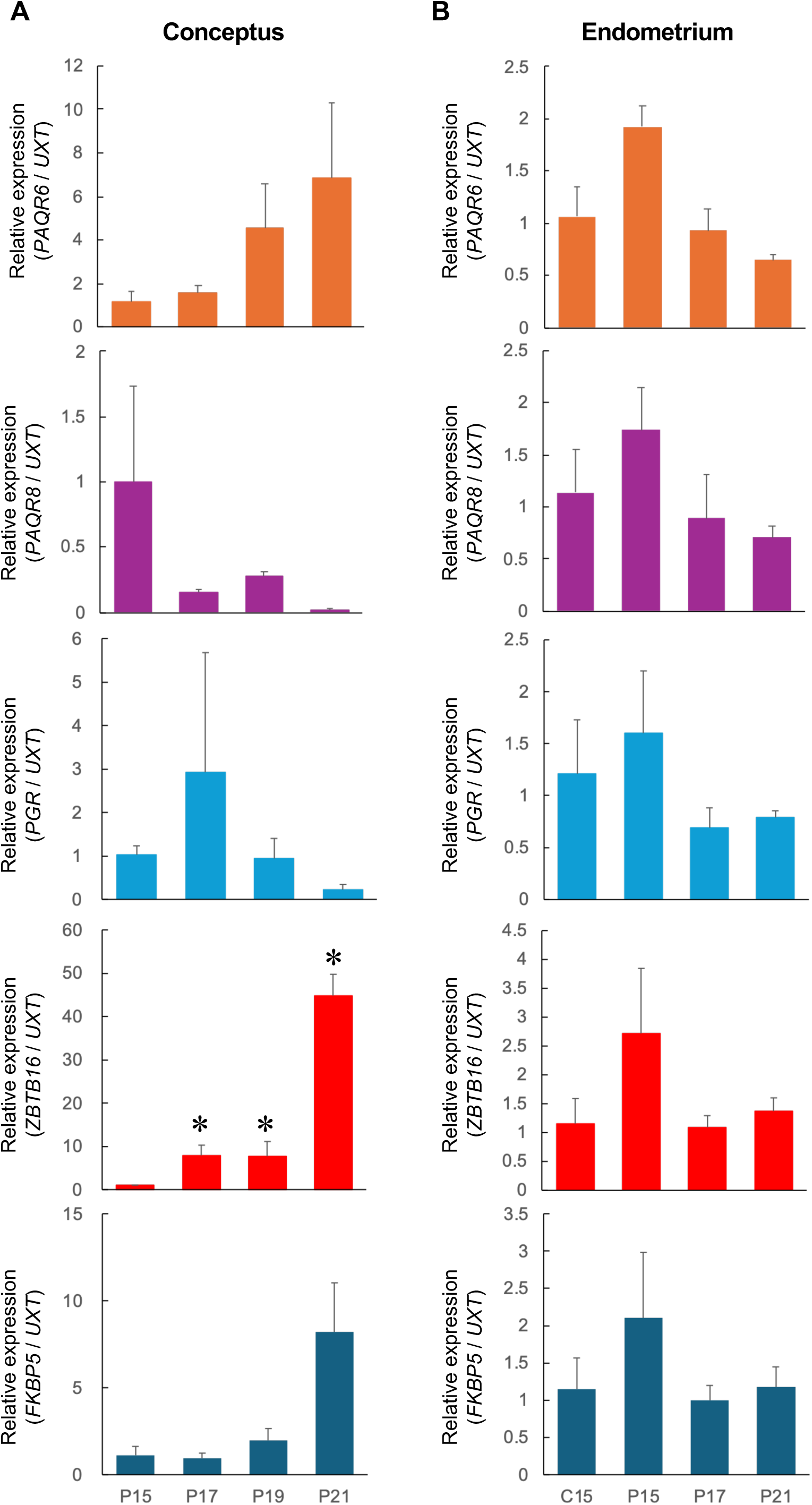
Transcript levels of PAQR6, PAQR8, PGR, ZBTB16 and FKBP5 in the ovine conceptus and endometrium across the peri-implantation period. Relative expression (target/ UXT , ΔΔCt method) by qPCR. (A) Conceptus at days 15, 17, 19 and 21 of pregnancy, normalized to day 15. (B) Endometrium at cyclic day 15 (C15) and days 15, 17 and 21 of pregnancy, normalized to C15. PAQR6 , PAQR8 and PGR did not change significantly in either tissue. *ZBTB16* increased significantly in the conceptus, whereas *FKBP5* showed a numerical increase without reaching statistical significance; neither gene changed significantly in the endometrium. Data are mean ± SEM, n = 3 per group (conceptus day 15, n = 2). p < 0.05 versus day 15 (one-way ANOVA on ΔCt, Dunnett’s test).

### PAQR6 and PAQR8 localize to the endometrial epithelium

At the protein level, PAQR6 and PAQR8 localized predominantly to the endometrial epithelium, both the luminal epithelium and the glandular epithelium, at cyclic day 15 and at days 15 and 21 of pregnancy, with weaker or negligible signal in the stroma (Fig. 3A, B). Both receptors were present in the epithelium across all stages examined, consistent with the stable transcript levels and in contrast to the stage-dependent loss of nuclear PGR reported for this epithelium.

**Fig. 3.**
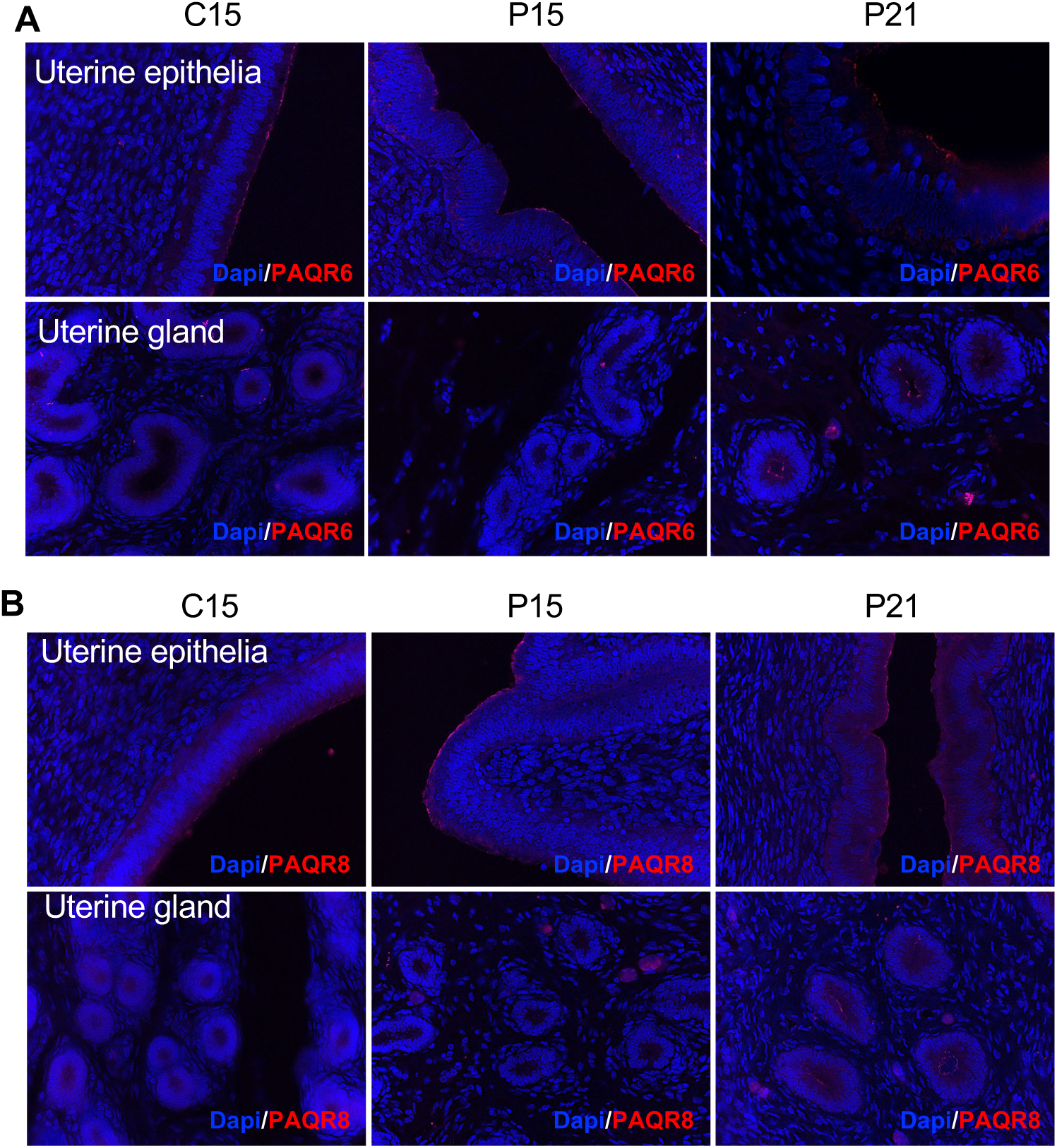
Localization of PAQR6 and PAQR8 in the ovine endometrium. Immunofluorescent localization of PAQR6 (A) and PAQR8 (B) in the luminal (uterine epithelia) and glandular (uterine gland) epithelium at cyclic day 15 (C15) and days 15 and 21 of pregnancy; nuclei counterstained with DAPI. Both receptors localized to the endometrial epithelium.

### PAQR6 and PAQR8 are enriched at the apical epithelial surface

To locate the epithelial signal relative to the cell membrane, we co-stained PAQR6 or PAQR8 with the basolateral adherens-junction marker E-cadherin in the ovine luminal epithelium at day 21 of pregnancy. Both receptors (red) were enriched as a band along the apical surface of the epithelium, with weaker signal towards the base, and neither overlapped the E-cadherin signal (green), which outlined the lateral cell borders (Fig. 4A, B). PAQR6 and PAQR8 therefore show a junction-excluded distribution enriched at or near the apical surface of the luminal epithelium.

**Fig. 4.**
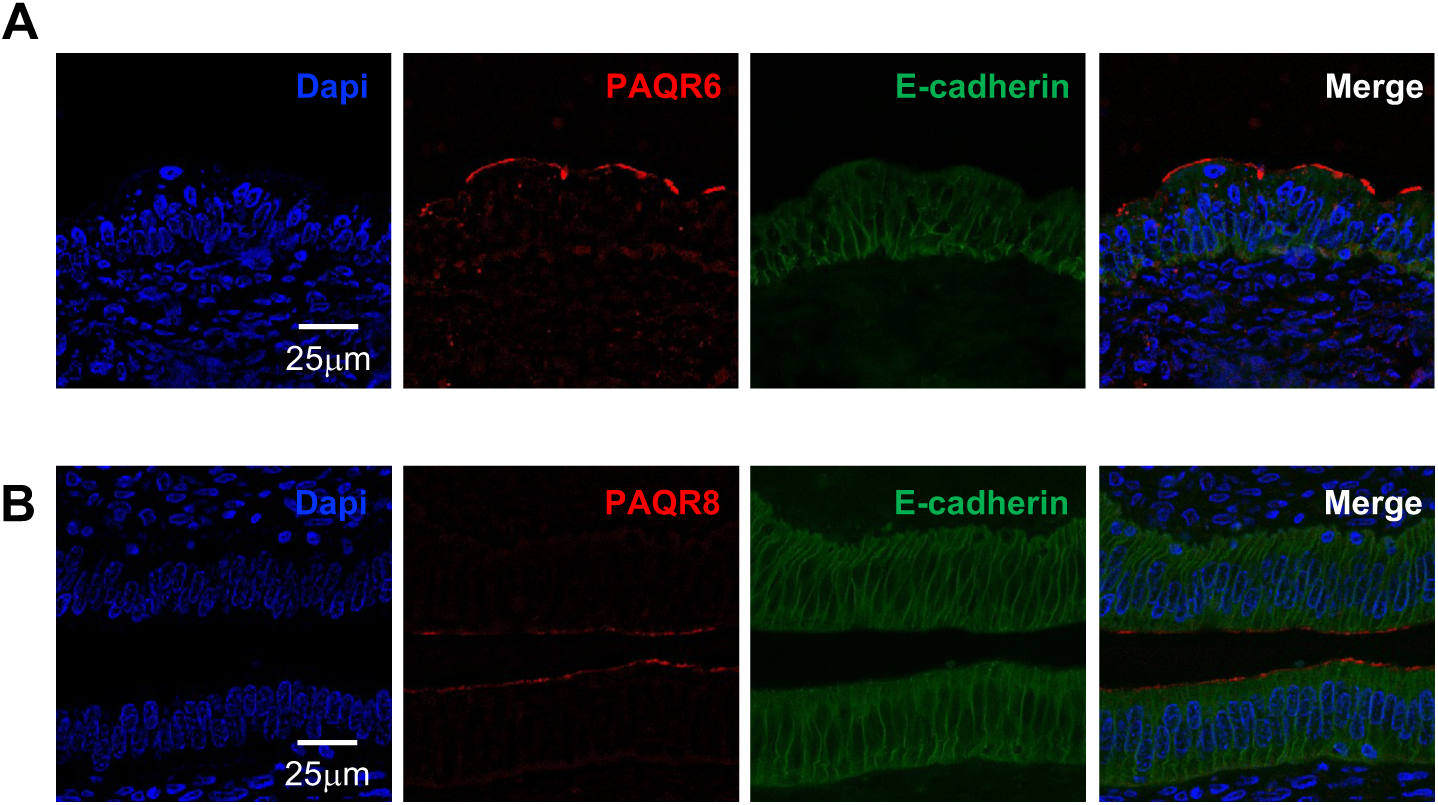
PAQR6 and PAQR8 are enriched at the apical surface of the ovine endometrial epithelium. Dual immunofluorescence of PAQR6 (A) or PAQR8 (B) (red) with the basolateral adherens-junction marker E-cadherin (green) in the ovine luminal epithelium at day 21 of pregnancy, with DAPI (blue) and merge. Both receptors were enriched at the apical surface and did not overlap E-cadherin. Scale bars, 25 µm.

### PAQR6 and PAQR8 localization in the bovine ovary

In the bovine ovary, PAQR6 and PAQR8 protein was detected in follicles across developmental stages (primordial, primary, secondary and antral/Graafian follicles) and in the corpus luteum, with signal in granulosa and luteal cells; preimmune serum gave no signal (negative control) (Fig. 5A, B).

**Fig. 5.**
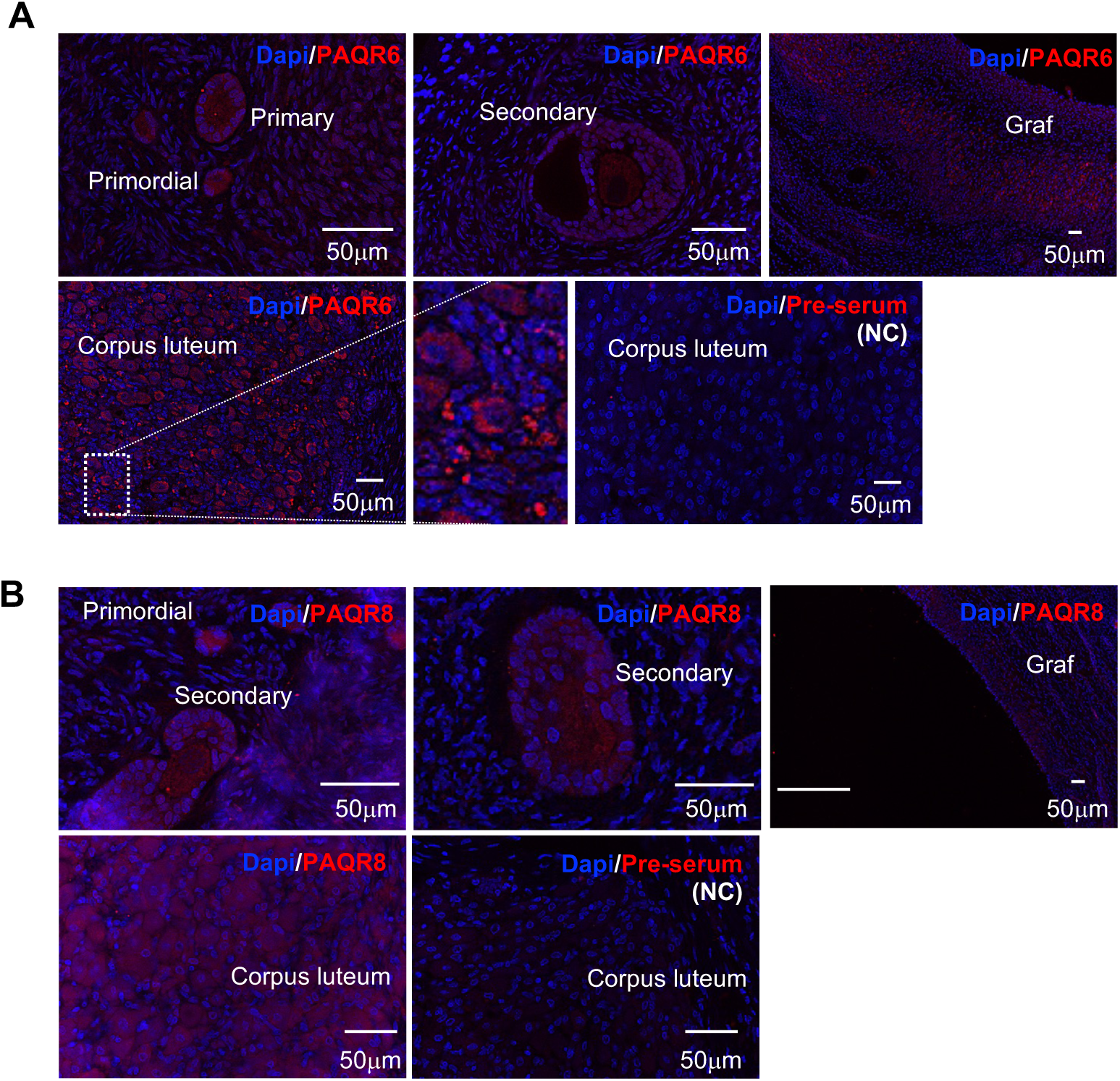
Localization of PAQR6 and PAQR8 in the bovine ovary. Immunofluorescent localization of PAQR6 (A) and PAQR8 (B) in follicles at successive developmental stages (primordial, primary, secondary, Graafian) and in the corpus luteum; nuclei counterstained with DAPI. Preimmune serum served as the negative control (NC). Scale bars, 50 µm.

## DISCUSSION

Progesterone action in the ruminant uterus has been examined almost entirely through the nuclear receptor PGR, and the membrane progesterone receptors that could mediate its rapid, PGR-independent effects had not been defined at the protein level in ruminant tissue, partly because antibodies validated in cattle or sheep were lacking. Using self-made, rigorously validated antibodies together with quantitative PCR, we provide the first combined transcript-and-protein map of the membrane progesterone receptors PAQR6 and PAQR8 in the ruminant uterus, conceptus and ovary. Two findings stand out: the dynamic genomic progesterone response over early pregnancy is confined to the conceptus, and PAQR6 and PAQR8 protein is retained in the endometrial epithelium throughout the peri-implantation period, where both receptors are enriched at the apical epithelial surface.

This study revealed a tissue-specific pattern of progesterone-responsive gene expression. In the conceptus, *ZBTB16* increased significantly as pregnancy advanced, reaching approximately 45-fold the day-15 level by day 21, whereas *FKBP5* showed a numerical increase that did not reach statistical significance. In contrast, *PGR*, *PAQR6* and *PAQR8* did not change significantly during this period. The stable whole-endometrial *PGR* transcript does not contradict the reported epithelial PGR down-regulation in sheep (Spencer and Bazer, 1995; Spencer et al., 2004), which is confined to the luminal and superficial glandular epithelium and would be masked in bulk-tissue qPCR by the PGR-retaining stroma and deep glands. These findings indicate that progesterone-responsive gene expression changes more dynamically in the conceptus than in the maternal endometrium during days 15-21 of pregnancy. However, because functional progesterone signaling was not directly assessed, these expression patterns should not be interpreted as direct evidence of increased genomic progesterone signaling.

Against this background, we examined the subcellular distribution of PAQR6 and PAQR8 in the endometrial epithelium. Co-staining placed both receptors at the apical surface of the ovine luminal epithelium, enriched apically and excluded from the E-cadherin-positive adherens junctions. The apical surface of the luminal and glandular epithelium faces the uterine lumen, so an apically enriched receptor is positioned to sense a luminal ligand directly, independently of the genomic axis, a pathway that could persist even as epithelial PGR is down-regulated at receptivity.

Apical localization of a signaling receptor in this epithelium has precedent. The membrane progesterone receptors mPRβ (PAQR8) and mPRγ localize to the apical, ciliated surface of the mammalian fallopian tube epithelium (Nutu et al., 2009), and cytokine receptors of the LIF/gp130 system act at the luminal epithelial surface during implantation (Namiki et al., 2023). Two caveats apply to our assignment. First, E-cadherin marks the lateral adherens junctions rather than the apical or basal membrane itself, so exclusion from E-cadherin together with apical morphology defines a junction-excluded, apically enriched distribution; a positive apical-versus-basal assignment will require apical (e.g. ezrin, acetylated tubulin) and basolateral (e.g. Na+/K+-ATPase) surface markers. Second, mPRs have in some systems been reported in intracellular or endoplasmic-reticulum compartments rather than at the plasma membrane (Krietsch et al., 2006), so distinguishing the apical plasma membrane from a sub-apical compartment will require higher-resolution imaging.

That both PAQR6 and PAQR8 are epithelial, rather than partitioning between epithelium and stroma, argues against a simple division of labor between the two subtypes in the endometrium and suggests they may act at the same membrane. In the ovary, both receptors were present in follicles and corpora lutea, placing them in the granulosa and luteal cells that are the principal ovarian targets of progesterone, consistent with the established role of mPRs in follicular and oocyte physiology (Zhu et al., 2003a; Dressing et al., 2011).

Returning to the endometrium, the identity of the ligand sensed at this apical surface is the central open question, and the localization itself argues against the simplest explanation, that progesterone is the sole ligand. Progesterone is the canonical mPR ligand and the obvious first candidate, yet two considerations make it an unsatisfying sole explanation. First, a cell that needs to respond to progesterone already possesses the high-affinity nuclear receptor PGR; retaining a distinct membrane receptor once epithelial PGR has been lost implies a role not wholly redundant with the genomic pathway: a different ligand, a different (rapid, membrane-delimited) mode of action, or a different source of ligand. Second, luteal progesterone reaches the endometrium hematogenously, through the sub-epithelial vasculature and stroma, and therefore approaches the epithelium from its basal side; a receptor dedicated to sensing systemic progesterone would be expected basolaterally rather than at the apical surface facing the lumen. Although progesterone is lipophilic and not strictly confined by cell polarity, the apical enrichment is more readily reconciled with a ligand presented from the luminal compartment than with blood-borne progesterone. The luminal fluid of the reproductive tract carries a spectrum of steroids and steroid metabolites, including reduced progesterone metabolites, as documented in the bovine oviduct (Lamy et al., 2016), and several mPRs, including PAQR6 (mPRδ), respond not only to progesterone but to such reduced metabolites and other neuroactive steroids (Pang et al., 2013). We therefore propose that apically enriched PAQR6 and PAQR8 may sense luminal steroid analogues rather than, or in addition to, progesterone itself, a possibility that would give physiological meaning to the retention of a membrane (mPR) receptor as the genomic (PGR) pathway is lost from the epithelium. This remains a hypothesis: neither the luminal steroid composition nor the ligand selectivity of the ruminant receptors was examined here.

Several limitations apply. The work is largely descriptive: it establishes where these receptors are and how the genomic axis is partitioned, not what the membrane receptors do. The antibodies were raised against bovine peptides and do not cross-react with mouse, so genetic knockout tissue could not be used as an additional specificity control; specificity rests instead on peptide pre-absorption, preimmune comparison and high ELISA titer, supported by the near-identity of the bovine and ovine epitopes (Fig. 1A). The uterine/conceptus and ovarian datasets come from different ruminant species (ovine and bovine) and are presented as parallel descriptions rather than a single cross-species claim. Finally, the membrane-domain analysis rests on exclusion from the E-cadherin-positive junctions together with apical morphology, so a definitive apical-versus-basal assignment awaits apical and basolateral surface markers.

Beyond these specific findings, the validated anti-PAQR6 and anti-PAQR8 antibodies may serve as a useful resource for ruminant reproductive biology. Raised against bovine C-terminal epitopes that are near-identical in sheep and confirmed by peptide pre-absorption and preimmune comparison, they recognize these two subtypes in ruminant tissue, for which validated mPR antibodies have so far been scarce. In addition to the in vivo mapping presented here, such antibodies should be applicable to isolated, cultured ruminant endometrial and ovarian cells, for example, in the search for candidate ligands and in functional analyses of PAQR6- and PAQR8-mediated, membrane-initiated progesterone signaling, addressing questions that whole-tissue surveys cannot resolve.

In summary, PAQR6 and PAQR8 are epithelial membrane progesterone receptors of the ruminant endometrium, enriched at the apical epithelial surface and excluded from the adherens junctions, and are expressed in follicular and luteal cells of the ovary, while the dynamic genomic progesterone response of early pregnancy is a conceptus-specific event. Together with the validated subtype-specific antibodies, these maps provide a foundation for testing membrane-initiated progesterone signaling in ruminant reproduction.

## DISCLOSURES

The authors declare that they have no known competing financial interests or personal relationships that could have appeared to influence the work reported in this paper.

Declaration of generative AI and AI-assisted technologies in the manuscript preparation process During the preparation of this work, the authors used Claude (Anthropic) to assist with language editing and with drafting and refining the text. After using this tool, the authors carefully reviewed and edited the content as needed and take full responsibility for the content of the published article.

## DATA AVAILABILITY

The data supporting the findings of this study are available from the corresponding author upon reasonable request.

## AUTHOR CONTRIBUTIONS

H.Y., Y.Y. and K.N. conceived and designed research; H.Y., Y.Y., M.Y., Y.M. and K.I. performed experiments; H.Y. and K.N. analyzed data; Y.Y. and K.N. interpreted results of experiments; K.N. prepared figures; H.Y. and K.N. drafted manuscript; K.I., F.O., H.O., E.K. and I.K. edited and revised manuscript; all authors approved the final version of the manuscript.

