## Supplementary Table S1 for "Membrane progesterone receptors PAQR6 and PAQR8 in the ruminant uterus and ovary"

**Supplementary Table S1.** Primer sequences used for quantitative PCR.

| **Gene** | **Forward primer (5′–3′)** | **Reverse primer (5′–3′)** | **Product size (bp)** |
| --- | --- | --- | --- |
| *UXT* | TGGCAGAAGCTCTCAAGTTCATT | CATGTGGATATGGGCCTTGAT | 104 |
| *PGR* | TAATCTGTGGGGATGAAGCA | CAGCACTTTCTAAGGCGACA | 181 |
| *PAQR6* | CACCATGCTCAGTCTCAAGC | GACGGTCTCGTTGGTCATCT | 154 |
| *PAQR8* | CCTCCATCACGTACCTCACC | GAGCCAGAAGTGCTCGTACC | 179 |
| *ZBTB16* | CACTCAGCGGGTGCCAAAG | TTCCCACACAGCAGACAGAAG | 131 |
| *FKBP5* | CAGATCTCCATGTGCCAGAAA | TTACTGGCCTCTTCCTTTGC | 113 |

UXT, reference gene; bp, base pairs.
